# Bacterial metatranscriptomes in wastewater can differentiate virally infected human populations

**DOI:** 10.1101/2022.02.23.481658

**Authors:** Rodolfo A Salido, Cameron Martino, Smruthi Karthikeyan, Shi Huang, Gibraan Rahman, Antonio Gonzalez, Livia S. Zaramela, Kristen L Beck, Shrikant Bhute, Kalen Cantrell, Anna Paola Carrieri, Sawyer Farmer, Niina Haiminen, Greg Humphrey, Ho-Cheol Kim, Laxmi Parida, Alex Richter, Yoshiki Vázquez-Baeza, Karsten Zengler, Austin D. Swafford, Andrew Bartko, Rob Knight

## Abstract

Monitoring wastewater samples at building-level resolution screens large populations for SARS-CoV-2, prioritizing testing and isolation efforts. Here we perform untargeted metatranscriptomics on virally-enriched wastewater samples from 10 locations on the UC San Diego campus, demonstrating that resulting bacterial taxonomic and functional profiles discriminate SARS-CoV-2 status even without direct detection of viral transcripts. Our proof-of-principle reveals emergent threats through changes in the human microbiome, suggesting new approaches for untargeted wastewater-based epidemiology.

## Body

Our past work deploying a highly spatially resolved, high-throughput wastewater monitoring system on a college campus (1) enabled collection and qPCR characterization of thousands of wastewater samples, identifying 85% of SARS-CoV-2 clinical cases (2), and also enabling genomic surveillance for emerging variants of concern by complete genome sequencing from extracted RNA (3). Wastewater-based epidemiology (WBE) provides additional advantages in that it is (i) non-invasive, (ii) cost-effective relative to individual clinical testing, (iii) does not require individuals to consent to clinical testing that is often reported to public health agencies, and (iv) can therefore benefit under-served populations (4–6). However, this WBE scheme is currently limited to pathogen detection and characterization through targeted qPCR and sequencing, and cannot detect agents of disease for which a screening test has not been developed.

Here we describe an untargeted community/population level disease monitoring strategy using metatranscriptomics, which leverages correlations in observable changes in wastewater microbiomes with human microbiome disruptions associated with disease state. SARS-CoV-2, like many pathogens, has been reported to cause systematic disruptions in the human gut microbiome (7–9), which is the principal human microbial input to wastewater (10). We employed this strategy to test whether information in the wastewater metatranscriptome could discriminate SARS-CoV-2 positive from negative wastewater samples (assessed by qPCR) as a proof-of-principle.

We present a high-throughput wastewater metatranscriptomics pipeline that lowers the accessibility to an otherwise cost-prohibitive sequencing method at scale through miniaturization, parallelization, and automation (11–12). (**Sup. Fig. S1**) Using this pipeline, we generated metatranscriptomics sequencing data for 313 virally-enriched (VE) wastewater samples collected from manholes servicing different residential buildings across a college campus, including isolation housing buildings (Manhole IDs: C6M095-C6M098), from Nov 23 2020 to January 7 2021. Sequencing reads were demultiplexed, trimmed, and quality filtered before being deposited in Qiita (13), where ribosomal reads were removed using SortMeRNA (14) using default processing recommendations; non-ribosomal reads were aligned to genomes or genes using Woltka (15) resulting in two different feature tables: taxonomic and functional (details in Materials and Methods).

Samples obtained from each manhole have a distinct microbiome signature, likely a composite of the individual microbiomes of the people contributing to each wastewater stream. Beta-diversity analyses of both metatranscriptomic feature tables (taxonomic and functional) measured by Aitchison distance and robust Aitchison principal component analysis (RPCA) (16) reveal that wastewater samples cluster primarily by manhole source (manhole_id) (**Fig. 1A**), with a stronger signal than SARS-CoV-2 detection status (**Fig. 1B**)(**Sup. Table ST1**). Wastewater samples separate according to SARS-CoV-2 status based on these bacterial profiles alone, but this signal is obscured in the RPCA ordination by the stronger manhole_id clustering effect. Taxonomic features provide better separation by both SARS-CoV-2 status and manhole_id than functional features (**Supp. Table ST1**), suggesting that microbial community membership rather than current functional gene expression is more strongly affected by infection.

**Figure 1:**
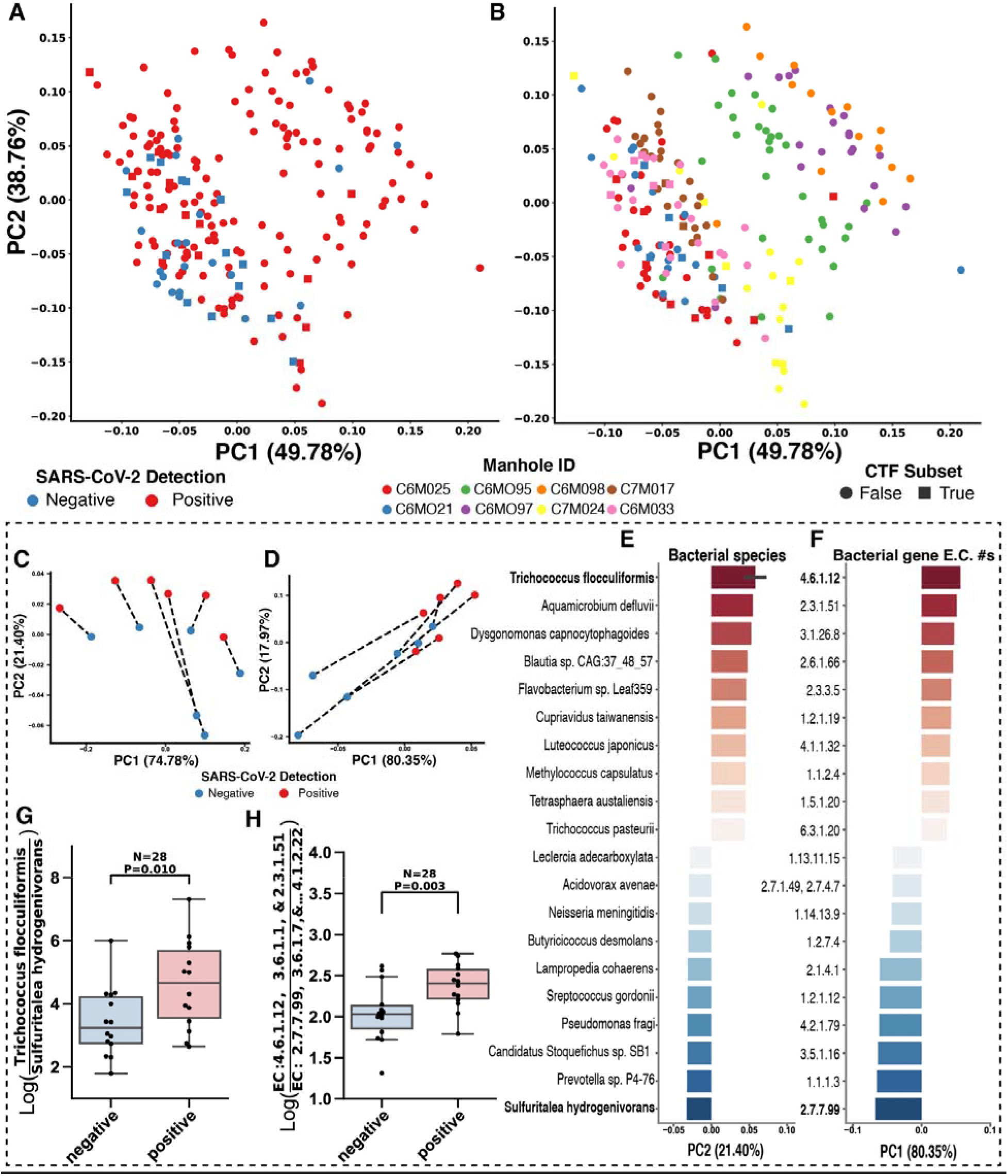
Microbial community composition changes can be observed in SARS-CoV-2 positive vs. negative wastewater samples. Robust principal component analysis (RPCA) of wastewater samples colored by SARS-CoV-2 detection status (**A**) and manhole source (**B**). A subset of samples (squares) was selected for pairwise comparisons of SARS-CoV-2 positive and negative wastewater microbiomes within a manhole and a week using compositional tensor factorization (CTF) on taxonomic (genomes, **C**) and functional (genes, **D)** features. Results shown in the dashed box are exclusive to this subset of samples. Important bacterial genomes (**E**) and genes (**F)** identified from CTF show significant differences between positive and negative sample groupings by log-ratios of top and bottom ranked features respectively (**G-H**). Error bar on the x-axis of the ranked features plot represents the standard error in the PC2 loadings across strains within the same species. The log-ratio boxplot elements are defined as follows: the centerline is the median of the distribution, box limits represent upper and lower quartiles, whiskers span 1.5x of the interquartile range, and points represent all data points.

To test whether the SARS-CoV-2 detection status-dependent microbiome signal can be identified even against the stronger manhole_id clustering effect, we selected a subset of samples for paired comparisons between SARS-CoV-2 positive and negative samples within specific manholes across one week (selection process detailed in Materials and Methods). This subset (squares, n=28 **Fig. 1A-B**) was analyzed by dimensionality reduction with compositional tensor factorization (CTF) (17), which accounts for the intra-manhole sample correlation. The resulting ordination shows that samples of the microbiome in any specific manhole undergo a pronounced shift along one of the main principal components (PC1 for taxonomic, PC2 for functional), when the subject population it services becomes infected with SARS-CoV-2 (**Fig. 1C-D**). Consequently, taxonomic features (genomes) that drive segregation along PC2 (**Fig. 1E**), or functional features (genes) along PC1 (**Fig. 1F**), can be positively or negatively correlated with SARS-CoV-2 detection. Log-ratio analysis of the top and bottom ranked taxonomic features as numerator and denominator respectively show a significant difference in the means of the SARS-CoV-2 detection sample groupings (**Fig. 1G**). Similarly, a log-ratio of six functional features positively and negatively ranked along PC2 also shows a significant difference in the means of the SARS-CoV-2 detection sample groupings (**Fig. 1H**) (see Materials and Methods).

The predictive power for wastewater SARS-CoV-2 status discrimination of the features selected through CTF analysis was validated via log-ratios and random forest machine learning (RFML) classification, using the remaining samples in this study (circles, **Fig. 1A-B**) plus an additional validation set (total n=285, positive=179, negative=106, **Sup. Table ST2**). Log-ratios of selected taxonomic and functional features showed a significant difference by SARS-CoV-2 detection status across the validation sampleset, with function (*t*-test, *T*=-3.9 p=0.0001) (**Fig. 2A**) showing a smaller effect than taxonomy (*t*-test, T=-8.8, p=1.3e-16) (**Fig. 2B**). Type II ANOVA of both log-ratios shows that differences in sample means are larger across SARS-CoV-2 status groups than manhole_id or sample_plate confounders (**Sup. Fig. S2**). The performances of the RFML classification models were evaluated through average area under the curve of precision-recall (AUC-PR) tests of stratified 5-fold cross validation classification tasks distinguishing samples’ SARS-CoV-2 status, manhole_id, and sample_plate. Lower dimensional feature tables from feature selection show comparable SARS-CoV-2 status classification performance as full feature tables for both data modalities (taxonomic and functional) (**Fig. 2C**), but reduced classification performance when distinguishing confounding manhole_id (**Fig. 2D**) or sample_plate (**Sup. Fig. S3**).

**Figure 2:**
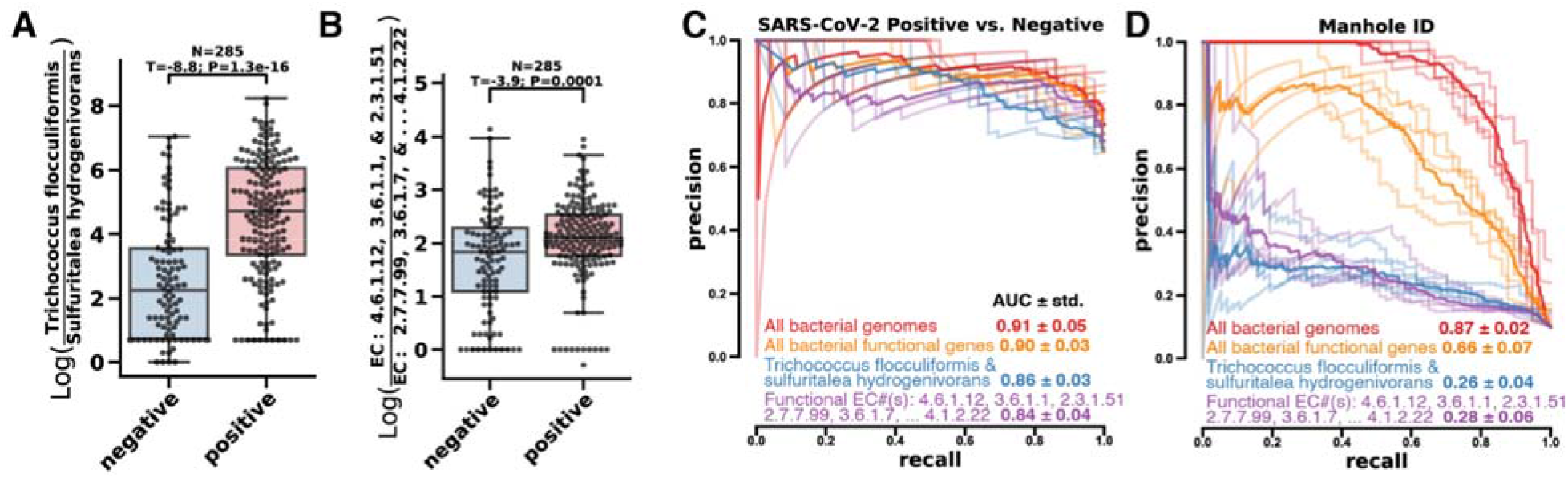
Key bacterial features identified in small paired subset show significant differences in larger validation dataset and provide RFML the ability to accurately predict SARS-CoV-2 status but not manhole source in wastewater. Log-ratios of important features, taxonomic (**A**) and functional (**B**), identified by CTF significantly separate wastewater samples by SARS-CoV-2 detection status in the remaining samples not included in the CTF subset. The log-ratio boxplot elements are defined as follows: the centerline is the median of the distribution, box limits represent upper and lower quartiles, whiskers span 1.5x of the interquartile range, and points represent all data points. **C**) Random forest machine learning 5-fold cross-validation shows high precision-recall of samples with positive SARS-CoV-2 detection status from taxonomic and functional tables with all features or a few selected features. **D**) Feature selection reduces Manhole ID classification performance while retaining SARS-CoV-2 discrimination, suggesting a reduction of overfitting. The translucent precision-recall curve traces of each feature table reflect all 5-fold cross-validation results while the bold trace represents the average.

Our results demonstrate that wastewater metatranscriptomes can reveal traces of rare pathogens through alterations of the microbiome of the afflicted individuals, which are eventually reflected in the wastewater microbiome. When effects are confounded by site/population, leveraging generalizable log-ratios separating positive/negative groupings across sites reduces overfitting. This proof-of-principle justifies further research on high-throughput wastewater metatranscriptome biomarker discovery for WBE; the untargeted nature of this data modality makes it flexible enough to monitor multiple diseases at the population scale (through traditional direct detection of known sequences from pathogens, but also by leveraging microbiome perturbations as a proxy), and is superior to metagenomic monitoring because it encompasses all living organisms and viruses(18). One of the limitations of the proposed strategy is the narrow stability of the samples’ RNA molecules. However, our methods don’t claim to comprehensively characterize the wastewater metatranscriptome and instead focus on the fact that changes in the observable bacterial metatranscriptome are sufficient to discriminate the wastewater’s viral status, with SARS-CoV-2 detection status serving as a relevant case study. Although key features of the bacterial metatranscriptome discriminate SARS-CoV-2 detection, further work is needed to determine how broadly this phenomenon generalizes to other pathogens. Lastly, our methodology allows automated high-throughput metatranscriptomics processing, applicable to many biospecimen types, and could have considerable impact beyond WBE.

## Acknowledgments

This work is supported in part by the IBM Research AI through the AI Horizons Network, IBM Artificial Intelligence for Healthy Living (A1770534), UC San Diego’s Return to Learn Program, NIH Director’s Pioneer Award (DP1AT010885), NSF RAPID Award (# 2038509), and Emerald Foundation Distinguished Investigator Award.

---

**Supplementary Figure S1:**
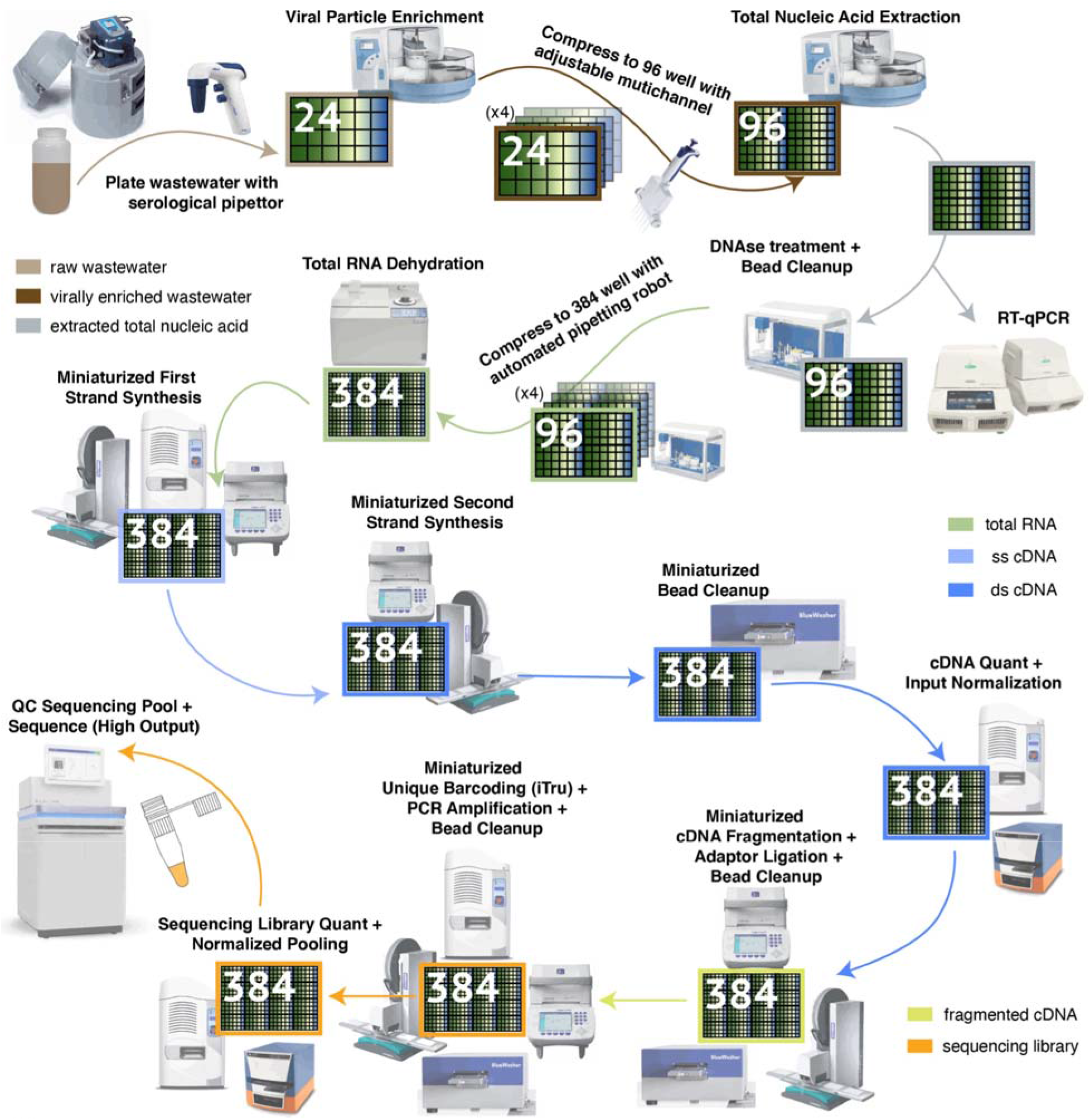
High Throughput pipeline for Virally Enriched (VE) wastewater metatranscriptomics. Flow diagram of metatranscriptomic data generation from VE wastewater samples, from auto-sampler to sequencer. Key robotic instrumentation and tools are depicted alongside each step. The flow diagram is color coded according to the different stages of sample processing. The high throughput pipeline increases sample processing parallelization through incremental compression of samples from 24-well plates to 384-well plates. Significant per sample cost savings are achieved through miniaturization of molecular reactions in 384-well format, for which specialized low volume liquid handling infrastructure is needed.

**Supplementary Table ST1:** PERMANOVA results on RPCA distance matrix show stronger manhole of origin effect than SARS-CoV-2 status. An analysis of variance of the Aitchison distance between wastewater samples shows that manhole of origin has the strongest effect size, followed by sample processing plate, and SARS-CoV-2 status. Samples from different manholes were not uniformly distributed across sample processing plates, confounding the effect sizes for both independent variables.

|  |  |  | PERMANOVA: F-stat. | PERMANOVA: p-value |
| --- | --- | --- | --- | --- |
| across-all | taxonomic | manhole_id | 23.9008 | 0.0002 |
|  |  | time_encoded | 0.8869 | 0.7321 |
|  |  | sars_cov_2_status | 8.8129 | 0.0002 |
|  |  | sample_plate | 21.2303 | 0.0002 |
|  | functional | manhole_id | 11.9542 | 0.0002 |
|  |  | time_encoded | 0.9860 | 0.5055 |
|  |  | sars_cov_2_status | 4.0365 | 0.0180 |
|  |  | sample_plate | 9.1532 | 0.0002 |

**Supplemental Table ST2:**
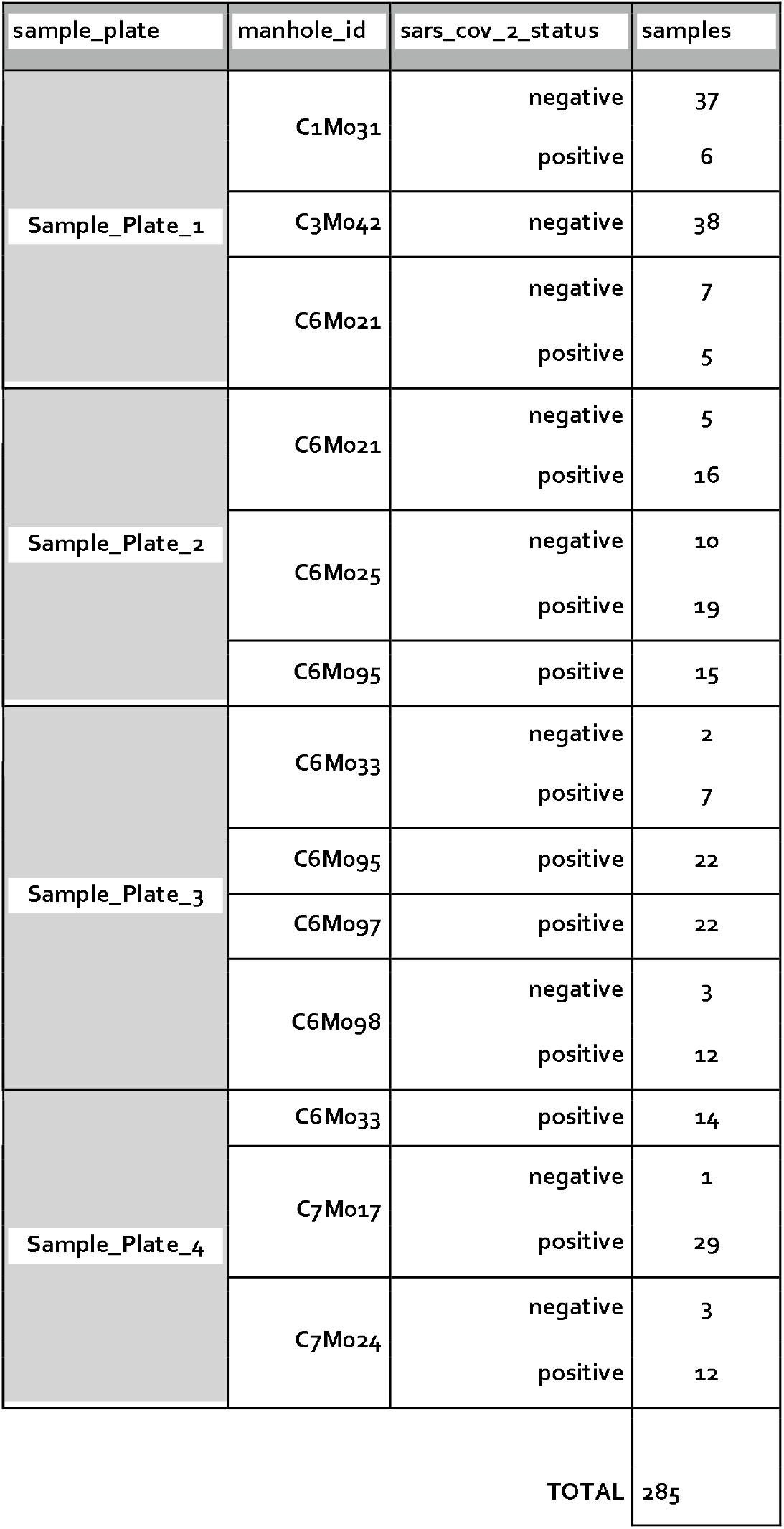
Description of validation dataset for Random Forest Machine Learning (RFML). Distribution of samples across different groupings relevant to the observed variance in the unsupervised learning analysis. Sample plate 1 was added, as an additional validation set, to the RFML analyses. The validation dataset excludes the subset of samples selected for the CTF analysis (n=28).

**Supplementary Figure S2:**
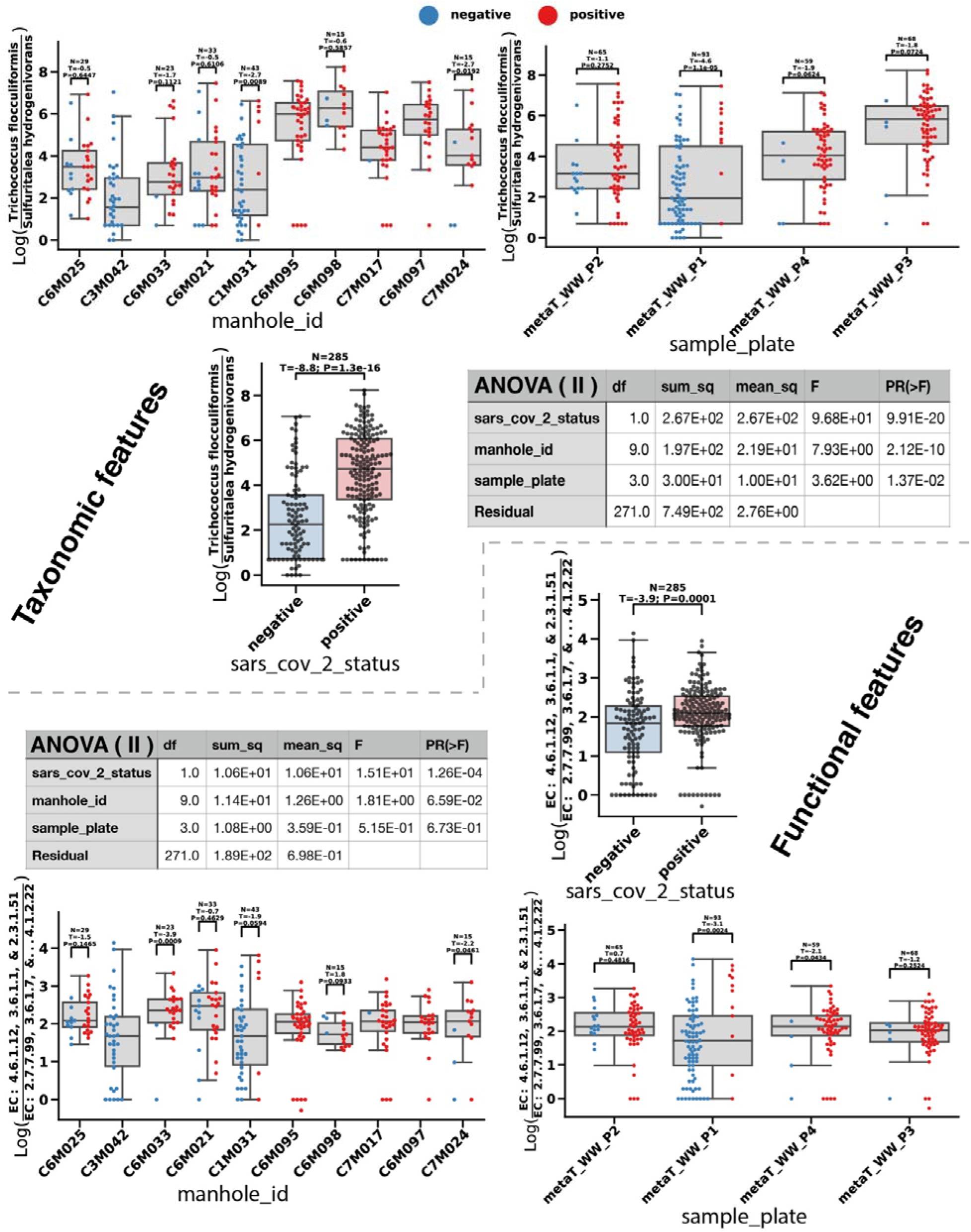
Analysis of variance (ANOVA) of both log-ratios show that SARS-CoV-2 status has the strongest effect size. Boxplots with overlaid swarmplots show the distribution of selected log-ratios for both taxonomic and functional feature tables, grouped by relevant sample metadata. The log-ratio boxplot elements are defined as follows: the centerline is the median of the distribution, box limits represent upper and lower quartiles, whiskers span 1.5x of the interquartile range, and points represent all data points. Results from ANOVA (type II) analyses are shown as tables for each feature modality. Statistical tests results (Student’s *t*-test) between SARS-CoV-2 status subgroupings (negative=blue / positive=red) in manhole_id and sample_plate plots are also shown, evidencing that the log-ratios generalize and perform better at discriminating SARS-CoV-2 status across all samples than within specific manholes.

**Supplementary Figure S3:**
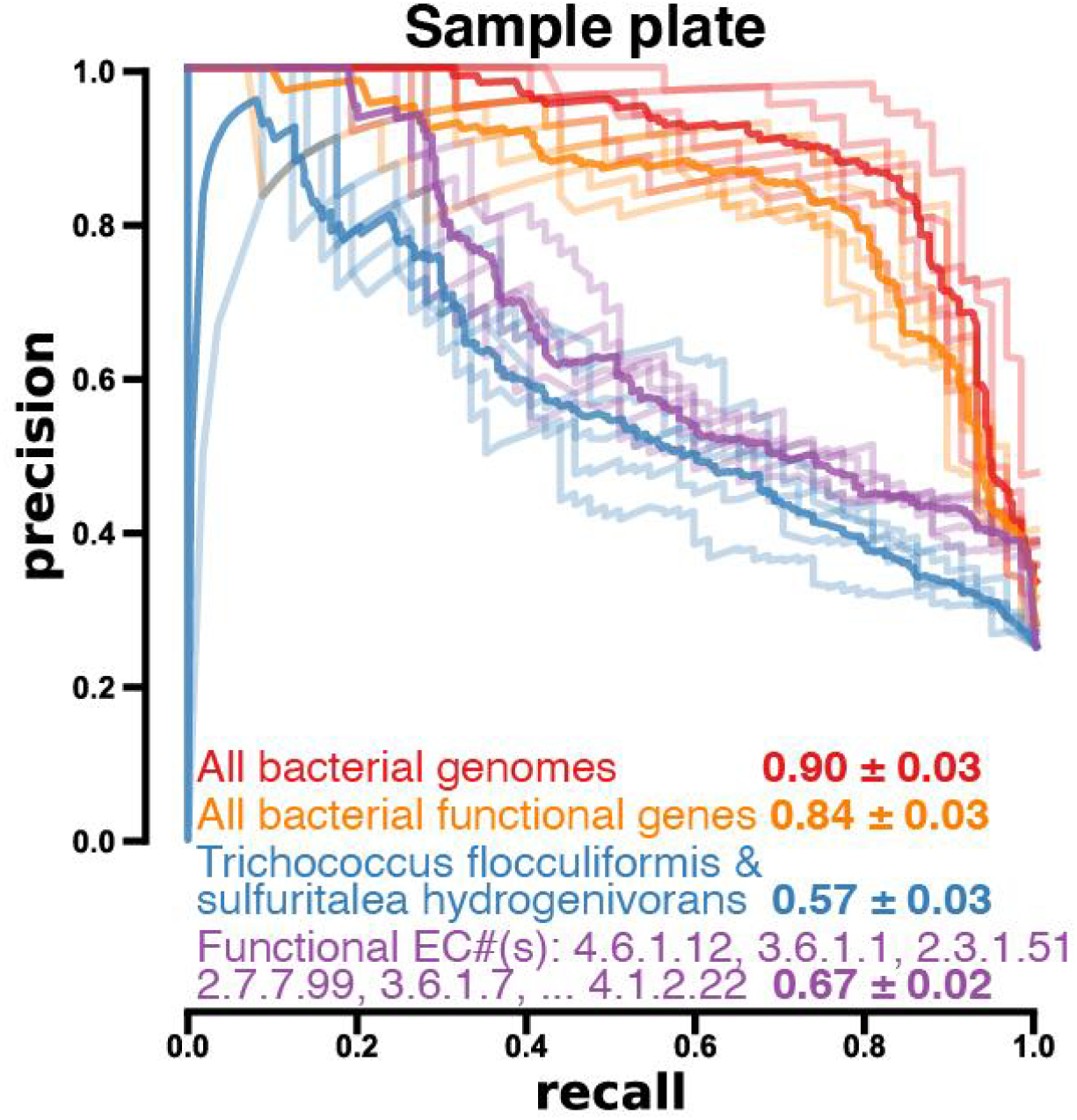
Random forest machine learning 5-fold cross-validation shows a decrease in precision-recall of samples’ processing plate (sample plate) from feature selection of taxonomic and functional feature tables in comparison to full feature tables, suggesting a reduction of overfitting on a possible technical confounder. The translucent precision-recall curve traces of each feature table reflect all 5-fold cross-validation results while the bold trace represents the average.

## References

1. Karthikeyan, S. et al. High-Throughput Wastewater SARS-CoV-2 Detection Enables Forecasting of Community Infection Dynamics in San Diego County. mSystems 6, (2021).

2. Karthikeyan, S. et al. Rapid, Large-Scale Wastewater Surveillance and Automated Reporting System Enable Early Detection of Nearly 85% of COVID-19 Cases on a University Campus. mSystems 6, 793–814 (2021).

3. Karthikeyan, S. et al. Wastewater sequencing uncovers early, cryptic SARS-CoV-2 variant transmission. medRxiv 2021.12.21.21268143 (2021). doi:10.1101/2021.12.21.21268143

4. Fielding-Miller, R. et al. Wastewater and surface monitoring to detect COVID-19 in elementary school settings: The Safer at School Early Alert project. medRxiv 2021.10.19.21265226 (2021). doi:10.1101/2021.10.19.21265226

5. Reitsma, M. B. et al. Racial/ethnic disparities in covid-19 exposure risk, testing, and cases at the subcounty level in California. Health Aff. 40, 870–878 (2021).

6. Lieberman-Cribbin, W., Tuminello, S., Flores, R. M. & Taioli, E. Disparities in COVID-19 Testing and Positivity in New York City. Am. J. Prev. Med. 59, 326–332 (2020).

7. Wu, Y. et al. Altered oral and gut microbiota and its association with SARS-CoV-2 viral load in COVID-19 patients during hospitalization. npj Biofilms Microbiomes 7, 61 (2021).

8. Xu, R. et al. Temporal association between human upper respiratory and gut bacterial microbiomes during the course of COVID-19 in adults. Commun. Biol. 2021 41 4, 1–11 (2021).

9. Gu, S. et al. Alterations of the Gut Microbiota in Patients With Coronavirus Disease 2019 or H1N1 Influenza. Clin. Infect. Dis. 71, 2669–2678 (2020).

10. Newton, R. J. et al. Sewage reflects the microbiomes of human populations. MBio 6, (2015).

11. Sboner, A., Mu, X. J., Greenbaum, D., Auerbach, R. K. & Gerstein, M. B. The real cost of sequencing: Higher than you think! Genome Biol. 12, 1–10 (2011).

12. Mayday, M. Y., Khan, L. M., Chow, E. D., Zinter, M. S. & DeRisi, J. L. Miniaturization and optimization of 384-well compatible RNA sequencing library preparation. PLoS One 14, e0206194 (2019).

13. Gonzalez, A. et al. Qiita: rapid, web-enabled microbiome meta-analysis. Nat. Methods 15, 796–798 (2018).

14. Kopylova, E., Noé, L. & Touzet, H. SortMeRNA: fast and accurate filtering of ribosomal RNAs in metatranscriptomic data. Bioinformatics 28, 3211–3217 (2012).

15. Zhu, Q. et al. OGUs enable effective, phylogeny-aware analysis of even shallow metagenome community structures. bioRxiv 2021.04.04.438427 (2021). doi:10.1101/2021.04.04.438427

16. Martino, C. et al. A Novel Sparse Compositional Technique Reveals Microbial Perturbations. mSystems 4, e00016–19 (2019).

17. Martino, C. et al. Context-aware dimensionality reduction deconvolutes gut microbial community dynamics. Nat. Biotechnol. 2020 392 39, 165–168 (2020).

18. Sims, N. & Kasprzyk-Hordern, B. Future perspectives of wastewater-based epidemiology: Monitoring infectious disease spread and resistance to the community level. Environ. Int. 139, 105689 (2020).

