## Supplementary material for "Bacterial metatranscriptomes in wastewater can differentiate virally infected human populations": Materials and Methods

**Virally Enriched (VE) Wastewater Metatranscriptomics Pipeline**

Viral enrichment and total nucleic acid extraction

Total nucleic acid extraction from composite wastewater samples with a magnetic bead based viral enrichment step was performed as previously described (<https://www.protocols.io/view/high-throughput-wastewater-sars-cov-2-detection-pi-bshvnb66>). Briefly, 5mL of composite wastewater sample is aliquoted into (2) replicate 24-deepwell plates (ThermoFisher, 95040480) pre-filled with 75 µL of Nanotrap Magnetic Virus Particles (Ceres Nano, #44202), and a 24-deep well elution plate is filled with 500µL of MagMAX™ Microbiome Lysis Solution (ThermoFisher, #A42361). The 24-deep well wastewater and elution plates undergo a viral enrichment protocol in a KingFisher Flex (ThermoFisher). VE wastewater is then compressed into a 96-deep well plate (ThermoFisher, #95040450) with a variable span multichannel pipette. The 96-deep well VE wastewater plate then undergoes a nucleic acid extraction with a MagMAX™ Viral/Pathogen II (MVP II) Nucleic Acid Isolation Kit (ThermoFisher, #A48383), according to manufacturer’s recommendations, in a KingFisher Flex (ThermoFisher), with the following modification: 450 µL of VE wastewater were used as input for the extraction.

SARS-CoV-2 RT-qPCR detection

SARS-CoV-2 detection was performed as previously described (<https://www.protocols.io/view/high-throughput-wastewater-sars-cov-2-detection-pi-bshvnb66>), using the Promega SARS-CoV-2 RT-qPCR Kit for Wastewater (Promega, #CS317402) in a CFX384 Touch (BioRad) according to manufacturer’s recommendations and CDC recommended qPCR cycling (Promega GoTaq® Probe 1-Step RT-qPCR System, <https://www.fda.gov/media/134922/download>), with the following modification: a 10 µL total reaction size was used instead of the recommended 20 µL. Briefly, 4 µL of extracted nucleic acid were combined with a mastermix composed of 5 µL of GoTaq WW Master Mix (2X), 0.2 µL of GoScript RT (50X), 0.5 µL of 20X Prime/Probe/IAC Mix, and 0.3 µL of Nuclease Free Water in a 384-well PCR plate (Eppendorf, #951020702) which was then subjected to RT-qPCR.

DNAse treatment

Total nucleic acid extractions underwent automated DNAse treatment in 96-well format in an epMotion 5075 (Eppendorf) using TURBO™ DNase (ThermoFisher, #AM2238) following manufacturers recommendations with the following modifications: a total reaction size of 30 µL with 25 µL nucleic acid input and 0.5 µL of TURBO DNAse enzyme was used instead of the recommended 50 µL reaction with diluted nucleic acid input and 1 µL of TURBO DNAse enzyme. Inactivation of TURBO DNase was achieved through an automated 2X AMPure RNAClean XP (Beckman, #A63987) bead cleanup with two 80% ethanol washes performed in an epMotion 5075 (Eppendorf)). The final elution volume of the bead cleanup was 20 µL, but only 15 µL were recovered during automated elution due to bead contamination issues while pipetting.

Miniaturized cDNA synthesis

Total RNA underwent miniaturized cDNA synthesis in 384-well format, adapted from full size reaction protocols previously described (1). 10 µL per sample of Total RNA output from prior DNAse treatment step were compressed into a 384-well PCR plate (Eppendorf, #951020702) in a fully interweaved pattern using an epMotion (Eppendorf) (similar to <https://www.protocols.io/view/extracted-gdna-plate-compression-ua4esgw>). Total RNA was fully dried using a SpeedVac (ThermoFisher), and then resuspended in 2 µL of Nuclease Free Water (NFW), added with the Mosquito HV (SPT Labtech). Following SuperScript™ IV manufacturer recommendations, each sample of resuspended total RNA was primed for first strand synthesis with a mastermix of 200 nL of dNTP mix (10 mM each) and 200 nL of random hexamer primer (50 ng/µL) (supplied in SuperScript™ IV First-Strand Synthesis System, ThermoFisher, #18091050), which was dispensed in a single 400 nL mastermix transfer using an Echo 550 acoustic liquid-handling robot (Labcyte Inc.). A first strand synthesis mastermix with 200 nL of DTT (100 mM), 200 nL ribonuclease inhibitor (20 U/µL), 200nL of Superscriptase IV (200 U/µL), 800 nL of 5X first strand buffer (all of which are supplies in the SuperScript IV kit), and 200 nL of actinomycin D (5 mg/mL in 5% DMSO in NFW, ThermoFisher) per sample was mixed (1.6µL total) and transferred to the primed, total RNA using the Mosquito HV. After following thermal cycling conditions specified by SuperScript™ IV manufacturer for first strand synthesis, a second strand synthesis mastermix with 3 µL of 5X second strand buffer (Invitrogen™, 10812014), 400nL of dNTPS (10 mM), 200 nL of E. coli RNse H (2 U/µL, Invitrogen™, 18021071), 100nL of E. coli DNA ligase (10 u/µL, Invitrogen™, 18052019), and 100nL of E. coli DNA polymerase (ThermoFisher, EP0042) was mixed and added to the first strand synthesis output as a single 3.8 µL transfer using the Mosquito HV. An additional 7.2µL NFW transfer was performed to backfill the second strand synthesis reaction to a 15µL total reaction size, which was incubated at 16 ºC for 2 hours. Lastly, a 1.6X bead cleanup (Beckman, #A63987) with two 80% ethanol washes was performed using the BlueWasher (BlueCatBio). The final elution volume was 10 µL, but only 9 µL were recovered with Mosquito HV due to dead volumes. The eluted cDNA was plated in a low dead volume Echo qualified source plate (Labcyte, LP0200), and quantified for library prep input normalization using a miniaturized PicoGreen assay (ThermoFisher, #P7589).

Miniaturized Library Preparation

Library preparation of cDNA for next generation sequencing was performed as described previously (2). Briefly, a high-throughput version of the Kapa HyperPlus (Roche, #KK8514) library prep kit, miniaturized to an approximate 1/10 reagent volume and optimized for automated liquid handling, was used. cDNA input was normalized to 5ng in 3.5µL using the Echo. A Mosquito HV was used for 1/10 scale enzymatic fragmentation, end-repair, and adapter ligation reactions. Sequencing adapters and unique sample indexing were based on the iTru protocol (3), in which short universal y-yoke stub adapters are ligated first to fragmented cDNA and then a sample-specific barcode is added in a subsequent PCR step. Sub-microliter transfers of stub adapters and complimentary iTru barcode primers were dispensed with the Echo. High throughput bead cleanups were performed with a BlueWasher (BlueCatBio). Amplified, barcoded libraries were quantified by a miniaturized PicoGreen assay (ThermoFisher, #P7589) and pooled in approximately equimolar ratios for sequencing in a NovaSeq S4 flowcell (Illumina).

Sequencing

Next Generation Sequencing was performed on the NovaSeq 6000 (Illumina) instrument with S4 flowcells and paired end 150-bp runs. A first pass sequencing run in one lane of an S4 was performed, followed by a second pass sequencing run in another lane of an S4 with a rebalanced sequencing pool, based on non-ribosomal read counts from the initial sequencing run, of the same sequencing libraries

**Bioinformatics**

All sample preparations’ raw sequences were demultiplexed independently using default Illumina tools. Adaptors and human reads were removed using fastp [4] and minimap2 [5]; the resulting reads were uploaded to Qiita as study 13546 and processed using its default metatranscriptomics processing pipeline; in short: adaptor sequences were removed via fastp [3], then we separated ribosomal and non-ribosomal reads using Sortmerna v2.1b [6]; then, the non-ribosomal reads were processed with Woltka v0.1.1 [7] against the Web of Life (WoL) [8] reference to produce per-genome (taxonomic) and per-gene (functional) feature tables. The WoL per-gene feature tables were translated to functional annotations using the protein to UNIREF [9] map and then to KO [10] map via the collapse command in Woltka. The resulting independent, per preparation feature tables were merged into a single, aggregate feature table per feature modality.

**Dimensionality Reduction**

The data tables were filtered such that any feature had a total sum across samples of at least ten total counts and samples at least 700,000 and 100,000 total counts for the taxonomy and functional gene mapped respectively. Dimensionality reduction and beta diversity analysis of the data was performed through RPCA through Gemelli (v. 0.0.7) [11]. In order to create a paired set of data between positive and negative SARS-CoV-2 detections within a manhole site, the data were binned by time intervals and the number of paired samples was calculated; the maximum number of paired samples was obtained at 7-day intervals (one week). Compositional Tensor Factorization (CTF) was then used to perform dimensionality reduction and beta diversity analysis between manhole site and SARS-CoV-2 detection pairs through Gemelli (v. 0.0.7) [12]. The resulting Aitchison distances from both RPCA and CTF were assessed through permutational multivariate analysis of variance (PERMANOVA) [<https://doi.org/10.1002/9781118445112.stat07841>] using scikit-bio (v. 0.5.5). Analysis of variance (ANOVA) was performed through statsmodels (v. 0.13.1). In the case of multiple comparisons across all metrics Bonferroni *p*-value correction was applied.

**Feature Selection and Log-Ratios**

In order to explore those genes or genomes that were associated with the sample groupings, we selected features that were ranked along the same axis and direction as SARS-CoV-2 detection segregation in the CTF ordination. To validate the feature rankings positively/negatively associated with SARS-CoV-2 detection, log-ratios were produced. In the case of the taxonomic data, two features, the top and bottom ranked were used in the numerator and denominator respectively. The sparsity of the functional feature table excludes many samples when only one feature is used in the numerator and denominator of the log-ratio, so a compendium of features was chosen and aggregated for both numerator and denominator. The top four ranked features (top and bottom two) in addition to an additional E.C. number within the same functional annotation group were aggregated in the numerator and denominator (E.C. numbers, numerator: [3.6.1.1, 2.3.1.51, 4.6.1.12], denominator: [4.1.2.9 4.1.2.22, 2.7.7.99, 3.6.1.7]). Comparisons between the paired samples in the log-ratio were assessed through a paired *t*-test via Scipy (v. 1.7.3) [13].

**Supervised Machine Learning**

In order to explore the generalizability of the dimensionality reduction findings, machine learning classification by SARS-CoV-2 detection and Manhole ID was performed on the data not used in the CTF analysis. Random forest (RF) classification was performed through scikit-learn (v. 0.24.2) [<https://jmlr.org/papers/v12/pedregosa11a.html>] with n_estimators set to 500 and all other hyperparameters set to default. The data was split by 5-fold stratified cross-validation and the RF model was evaluated through the mean and standard deviation of the area under the precision recall curves for each fold of validation. This was repeated on each table and on the tables subset with features selected from the CTF analysis. The features selected in the CTF analysis were also compared by log-ratio (see previous section) between SARS-CoV-2 detection and manhole IDs through ANOVA and two-sided unpaired *t*-tests via Scipy (v. 1.7.3).

**References**

1. Embree, M., Nagarajan, H., Movahedi, N., Chitsaz, H. & Zengler, K. Single-cell genome and metatranscriptome sequencing reveal metabolic interactions of an alkane-degrading methanogenic community. *ISME J. 2014 84* **8**, 757–767 (2013).

2. Sanders, J. G. *et al.* Optimizing sequencing protocols for leaderboard metagenomics by combining long and short reads. doi:10.1186/s13059-019-1834-9

3. Glenn, T. C. *et al.* Adapterama I: Universal stubs and primers for 384 unique dual-indexed or 147,456 combinatorially-indexed Illumina libraries (iTru & iNext). *PeerJ* **2019**, e7755 (2019).

4. Chen, S., Zhou, Y., Chen, Y. & Gu, J. fastp: an ultra-fast all-in-one FASTQ preprocessor. *Bioinformatics* **34**, i884–i890 (2018).

5. Li, H. Minimap2: pairwise alignment for nucleotide sequences. *Bioinformatics* **34**, 3094–3100 (2018).

6. Kopylova, E., Noé, L. & Touzet, H. SortMeRNA: fast and accurate filtering of ribosomal RNAs in metatranscriptomic data. *Bioinformatics* **28**, 3211–3217 (2012).

7. Zhu, Q. *et al.* OGUs enable effective, phylogeny-aware analysis of even shallow metagenome community structures. *bioRxiv* 2021.04.04.438427 (2021). doi:10.1101/2021.04.04.438427

8. Zhu, Q. *et al.* Phylogenomics of 10,575 genomes reveals evolutionary proximity between domains Bacteria and Archaea. *Nat. Commun.* **10**, (2019).

9. Suzek, B. E., Huang, H., McGarvey, P., Mazumder, R. & Wu, C. H. UniRef: comprehensive and non-redundant UniProt reference clusters. *Bioinformatics* **23**, 1282–1288 (2007).

10. Kanehisa, M., Sato, Y., Kawashima, M., Furumichi, M. & Tanabe, M. KEGG as a reference resource for gene and protein annotation. *Nucleic Acids Res.* **44**, D457–D462 (2016).

11. Martino, C. *et al.* A Novel Sparse Compositional Technique Reveals Microbial Perturbations. *mSystems* **4**, e00016-19 (2019).

12. Martino, C. *et al.* Context-aware dimensionality reduction deconvolutes gut microbial community dynamics. *Nat. Biotechnol. 2020 392* **39**, 165–168 (2020).

13. Virtanen, P. *et al.* SciPy 1.0: fundamental algorithms for scientific computing in Python. *Nat. Methods* **17**, 261–272 (2020).

14. Hunter, J. D. Matplotlib: A 2D graphics environment. *Comput. Sci. Eng.* **9**, 90–95 (2007).
